# Predation increases prey fitness via transgenerational priming

**DOI:** 10.1101/2022.03.10.483883

**Authors:** Silvia Kost, Linea Katharina Muhsal, Christian Kost

## Abstract

Preparing your offspring for future challenges via priming can directly enhance its fitness. However, evidence for transgenerational priming has been limited to eukaryotic organisms. Here we test the hypothesis that predation primes bacteria such that their future generations respond with a more effective defence induction. In an evolution experiment, *Escherichia coli* was cultivated either in monoculture or in coculture with the predatory ciliate *Tetrahymena thermophila.* After 18 days, fitness and defensive clustering capabilities of derived bacterial populations were determined. Our results reveal that (i) predation can prime *E.coli* to induce their defensive cluster formation across generations and that (ii) three days of predation are sufficient to increase the fitness of predator-exposed over that of predator-free populations. Thus, our study shows that predation can have priming effects in bacterial populations that operate across generations, which concurs with the emerging perception that bacteria feature mechanisms to actively shape their evolutionary fate.

## INTRODUCTION

Predation is not only detrimental for the respective prey individual, but it also drives the dynamics of prey populations (Stevens, 2012). Due to its strong antagonistic effects, predation has significant consequences for the evolutionary fate of prey populations, acting as a potential driver for phenotypic divergence and speciation (Langerhans, 2007). To mitigate the negative effects of predation, prey organisms respond by avoiding, tolerating, or defending themselves against predators. For example, several animals reduce their activity and decrease their exploratory tendencies to escape predation (Lawler, 1989; Abbey-Lee *et al.*, 2016), while some plant seeds are able to tolerate herbivore attack due to their increased size (Xiao *et al.*, 2007). Defence mechanisms against predation have been intensively studied in plants, where the attack by herbivores leads to the induction of a multitude of responses (Karban & Baldwin, 1997). These so-called *induced defences* represent a form of phenotypic plasticity and it has been suggested that they have evolved to save the cost of constitutive defences in the absence of predation (Cipollini & Heil, 2010).

Induced plant defences can be primed, where upon a first (priming) stimulus the plant prepares itself for a subsequent (triggering) stimulus of the same kind, which ultimately allows for a faster and / or stronger defence response (Heil & Kost, 2006; Hilker *et al.*, 2016; Martinez-Medina *et al.*, 2016; Hilker & Schmülling, 2019). Even though the theory of plant defence priming mainly focuses on the effects predation has on certain individuals within the same generation, also transgenerational priming has been reported. In *Arabidopsis* for example, the F2 progeny of plants treated with *Pseudomonas syringae* were primed to activate inducible defence genes to counter future attack by this bacterial pathogen (Luna *et al.*, 2012).

Besides priming, prey populations can also show phenotypic plasticity across generations. In the context of predator-prey interactions, the term *transgenerational plasticity* (TGP) describes the phenomenon where the phenotype of a given generation is influenced by the presence of predators in previous generation(s), even if the current generation is not exposed to predation itself (Tariel *et al.*, 2020). Here, a famous example is the helmet-shaped heads of the water flea *Daphnia cucullata* (Agrawal *et al*., 1999): predator-exposed *Daphnia* do not only produce protective helmets themselves (i.e. within-generation plasticity), but also produce offspring that possesses the same defensive morphology (i.e. transgenerational plasticity).

However, not only eukaryotic organisms, but also microbial systems, have been shown to respond to predation. In these cases, a single predator frequently has the ability to engulf multiple cells of prey bacteria at a time. To mitigate negative fitness consequences, bacteria have evolved an array of adaptive responses against predation such as an increased swimming speed, the formation of multicellular clusters, or the production of toxins (Matz & Kjelleberg, 2005; Jousset, 2012). Over past decades, cluster formation has increasingly received attention, because it represents an induced defence mechanism that is performed by a cooperating group of bacteria. Specifically, individual partners invest into the production of costly structures that are needed to form multicellular groups (Dragoš *et al.*, 2018). As a consequence, cells may lose their Darwinian autonomy, thus making the predator-induced cluster formation relevant to understand the evolution of early multicellularity (Boraas *et al.*, 1998; Fischer *et al.*, 2016; Herron *et al.*, 2019). Regarding the defensive nature of clusters, these and other studies (Matz *et al.*, 2005) could show that the induced formation of cellular aggregates can increase resistance to predation, because larger bacterial aggregates exceed the size of the predator’s oral cavity. In terms of microbial predator-prey interactions, theoretical models emphasized the role of evolutionary rather than simple ecological factors shaping the underlying dynamics (Kaitala *et al.*, 2020). Especially because bacteria are typically characterized by very short generation times, predation may not only affect the current population, but is likely to also impact future generations. Thus, a key question in this context is: Does predation prime the induced defence responses of bacteria across generations?

While (transgenerational) priming and plasticity have not been described in a microbiological context before, bacteria and yeast possess two different mechanisms that enable them to prepare for future challenges: cross-protection and anticipation. *Cross-protection* refers to the situation in which one environmental stressor protects cells against another detrimental condition in the future. Examples involve starvation-induced protection against heat or H_2_O_2_ in *Escherichia coli* (Jenkins *et al.*, 1988) or oxidative stress-induced protection against salt stress in *Saccharomyces cerevisiae* (Dhar *et al.,* 2013). In the case of *anticipation*, current environmental conditions provide cues that signal future environmental change, which cells use to pre-emptively prepare for the corresponding conditions. A prominent example can be found in *E. coli*, which, when exposed to high temperatures, expresses genes that are adaptive when oxygen-levels drop. Interestingly, these are exactly the conditions *E. coli* faces when being orally ingested by a new host: temperature increases right after ingestion, which is followed by a decrease in ambient oxygen levels as cells enter the gastrointestinal tract (Tagkopoulos *et al.*, 2008).

Given that bacteria (i) form clusters as a powerful defence response to predation and (ii) are able to prepare for future challenges if they experience stressful conditions, we hypothesized that predation in bacteria should also induce transgenerational effects to enhance the fitness of future generations. To test this hypothesis, we serially propagated *E. coli* either in monoculture or in coculture with the predatory ciliate *Tetrahymena thermophila* (Figure 1). After an evolutionary period of 18 days, we determined fitness and clustering capability of the ancestral strain as well as of the mono- and the coevolved offspring either in the absence or in the presence of predators (i.e. naïve setting). Additionally, the fitness and clustering capability of mono- and coevolved offspring was compared after three days in the absence or presence of predators (i.e. experienced setting). Our results demonstrate that (i) predation can prime bacteria to induce defence responses across generations and that (ii) three days of predation are sufficient to increase the fitness of predator-exposed populations over that of predator-free populations.

**Figure 1.**
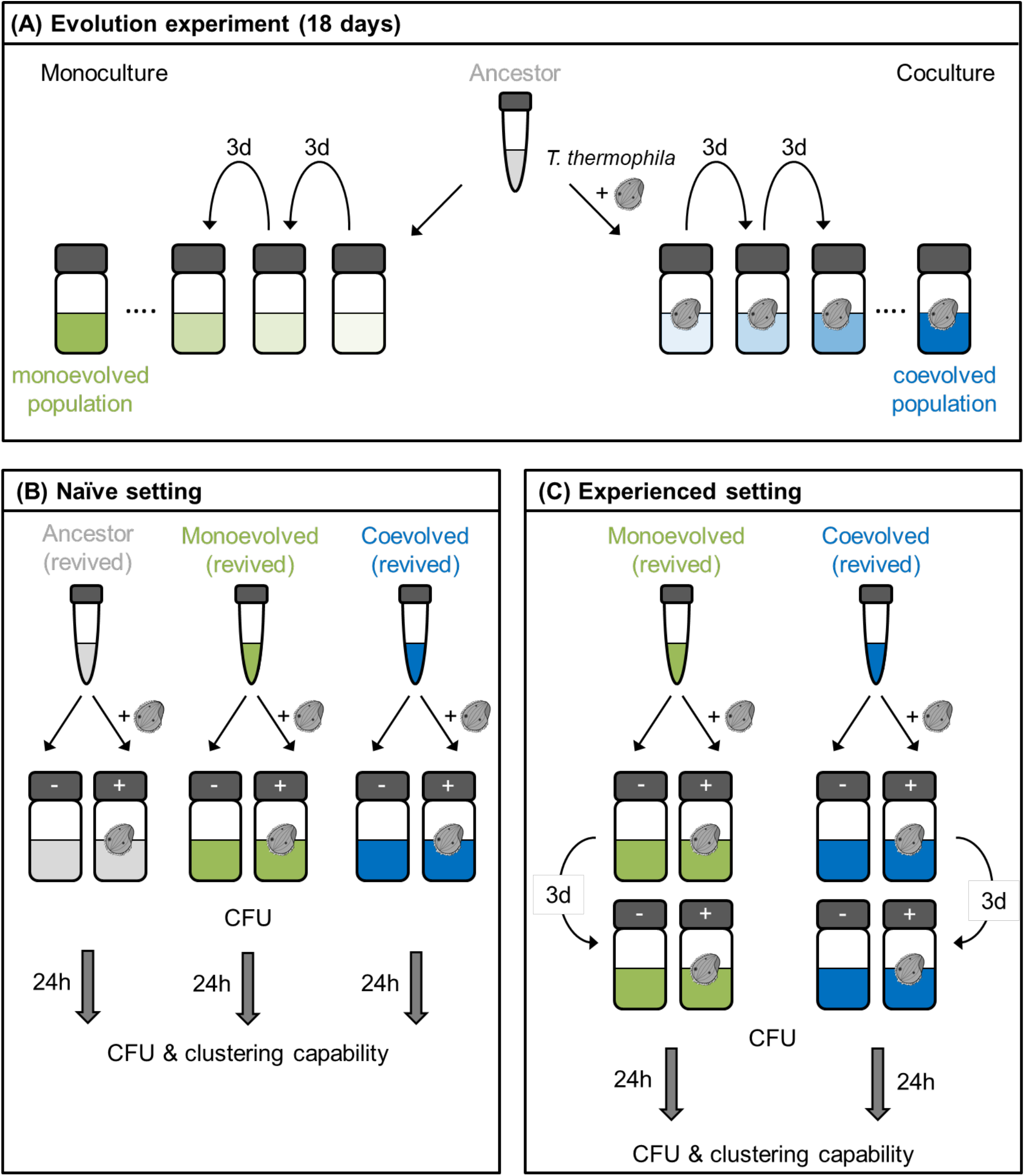
Overview over experimental conditions. **(A)** In the evolution experiment, populations of *E. coli* were either grown and serially propagated for a total of 18 days as monoculture or in coculture with the ciliate predator *T. thermophila.* Both treatment groups were initiated from one ancestral preculture and replicated four times. Cryostocks of the ancestral strain (grey) as well as of the mono - and coevolved offspring (green and blue, respectively) were stored at −80°C. **(B)** The naïve setting started with precultures of all three experimental groups. Given that *Tetrahymena* does not survive the freezing process, all freshly-started precultures were predator-free. From each preculture, one monoculture and one coculture were initiated and grown for 24 hours. At the beginning and at the end of the growth period, colony forming units (CFU) were counted. Clustering capability, expressed as the number of living cells per cluster, was evaluated after 24 hours. **(C)** The experienced setting comprised the mono- and coevolved offspring and resembled the naïve setting with the exception that experimental cultures were grown for three days and transferred once before measuring the CFU and clustering capability.

## MATERIAL AND METHODS

### Strains and culture conditions

*Escherichia coli* BW25113 was cultured in minimal medium for *Azospirillium brasilense* (MMAB) modified after Vanstockem et al. (1987) without biotin and supplemented with 0.5% of glucose (for precultures) or 0.05% of glucose (for experimental cultures) instead of malate. The predatory ciliate *Tetrahymena thermophila* (TSC_SD00026, C3 368.1, *Tetrahymena* stock center, Cornell University) was precultured axenically in proteose peptone medium (Cassidy-Hanley 2013) and subsequently transferred to MMAB with 0.05% of glucose for starvation and acclimatization to the experimental conditions. After 24 hours, starved cells were washed once with fresh MMAB containing 0.05% of glucose and immediately used in the respective experiments. All cultures were shaken continuously at 200 rpm and 30 °C.

### Evolution experiment

To test whether the exposure to predators induces transgenerational priming or plasticity of bacterial defence responses, *E. coli* was cultivated for 18 days in either the absence or the presence of *Tetrahymena thermophila* (Figure 1). The evolution experiment comprised four monoculture and four coculture replicate lines. All experimental lines were initiated from four individual parental precultures and set up in 100 ml medium with 0.005 bacterial OD_600nm_ (FilterMax F5 multi-Mode microplate reader, Molecular Devices). Cocultures were complemented with 638 *T. thermophila* cells per ml. Every three days, 5% of each culture was individually transferred into fresh medium. Cryostocks of the ancestral strain as well as of the mono- and coevolved offspring were stored at −80 °C.

### Coculture experiment: Naïve setting

Bacterial precultures of the ancestral strain and both evolved offspring types were initiated from their respective cryostocks (Figure 1). Please note that *Tetrahymena* does not survive the freezing procedure (Scheuerl *et al.*, 2019). Thus, all freshly started precultures were predator-free. From each preculture, two experimental cultures were set up as previously done in the evolution experiment in 100 ml medium with 0.005 bacterial OD_600nm_. Cocultures were complemented with 638 *T. thermophila* cells per ml. To calculate the Malthusian parameter as a measure of fitness according to Lenski *et al.* (1991), colony-forming units (CFU) were counted on MMAB agar plates at the beginning and at the end of a 24-hour growth period. To evaluate the cluster-forming capability of experimental cultures, aliquots of all populations were stained after growing for 24 hours using a bacterial viability kit (LIVE / DEAD^TM^ BacLight^TM^, Thermo Fisher). Samples were visualized with a confocal microscope (Zeiss LSM 880) and Z-stacks of three to five bacterial clusters per culture were recorded. Cell numbers in clusters were imaged and quantified using the Imaris software package (Oxford Instruments).

### Coculture experiment: Experienced setting

The experienced setting of the coculture experiment (Figure 1) was identical to the naïve one described above, yet with two exceptions. First, the experimental cultures were grown for three days and 5% of each culture was individually transferred once to fresh medium. Bacterial fitness and capability to form clusters were assessed at the beginning (i.e. 0 h) and at the end of the subsequent growth period (i.e. after 24 h) as before. Second, since the monoevolved offspring and the ancestral strain are redundant with respect to their predation experience, only the mono- and coevolved offspring were compared in the experienced setting.

### Statistical analysis

Data were analysed using the software package IBM® SPSS® Statistics 25. To meet test assumptions (i.e. homogeneity of variances and normal distribution), data were transformed if necessary.

## RESULTS

### Predation induces the adaptive formation of multicellular clusters across generations

To test whether the exposure to predators induces transgenerational defence responses in bacteria, *E. coli* was subjected to an evolution experiment, which was followed by two parallel coculture experiments (Figure 1). After the evolution experiment, the performance of the ancestral strain as well as of the mono- and coevolved offspring was analysed under both naïve (Figure 1B) and experienced conditions (Figure 1C).

In the naïve setting (Figure 2A), the fitness of the ancestral strain was significantly reduced in the presence of predation compared to the no-predator control (ANOVA followed by Tamhane’s T2 posthoc test of ln-transformed, absolute data: P < 0.001, n = 4). The same result was observed in offspring of monoevolved populations, where the fitness of all replicate populations was significantly lower in the presence of the predatory ciliate as compared to the no-predator control (ANOVA followed by Tamhane’s T2 posthoc test of ln-transformed, absolute data: P = 0.002, n = 8, Figure 2A). Remarkably, offspring of coevolved populations did not suffer from a significant fitness reduction in the presence of predation compared to the no-predation scenario (ANOVA followed by Tamhane’s T2 posthoc test of ln-transformed, absolute data, p = 1.0, n = 8, Figure 2A). In contrast, fitness of both types of coevolved populations (i.e. with and without predation) remained consistently high and on the same level as predator-free, monoevolved populations (Figure 2A).

**Figure 2.**
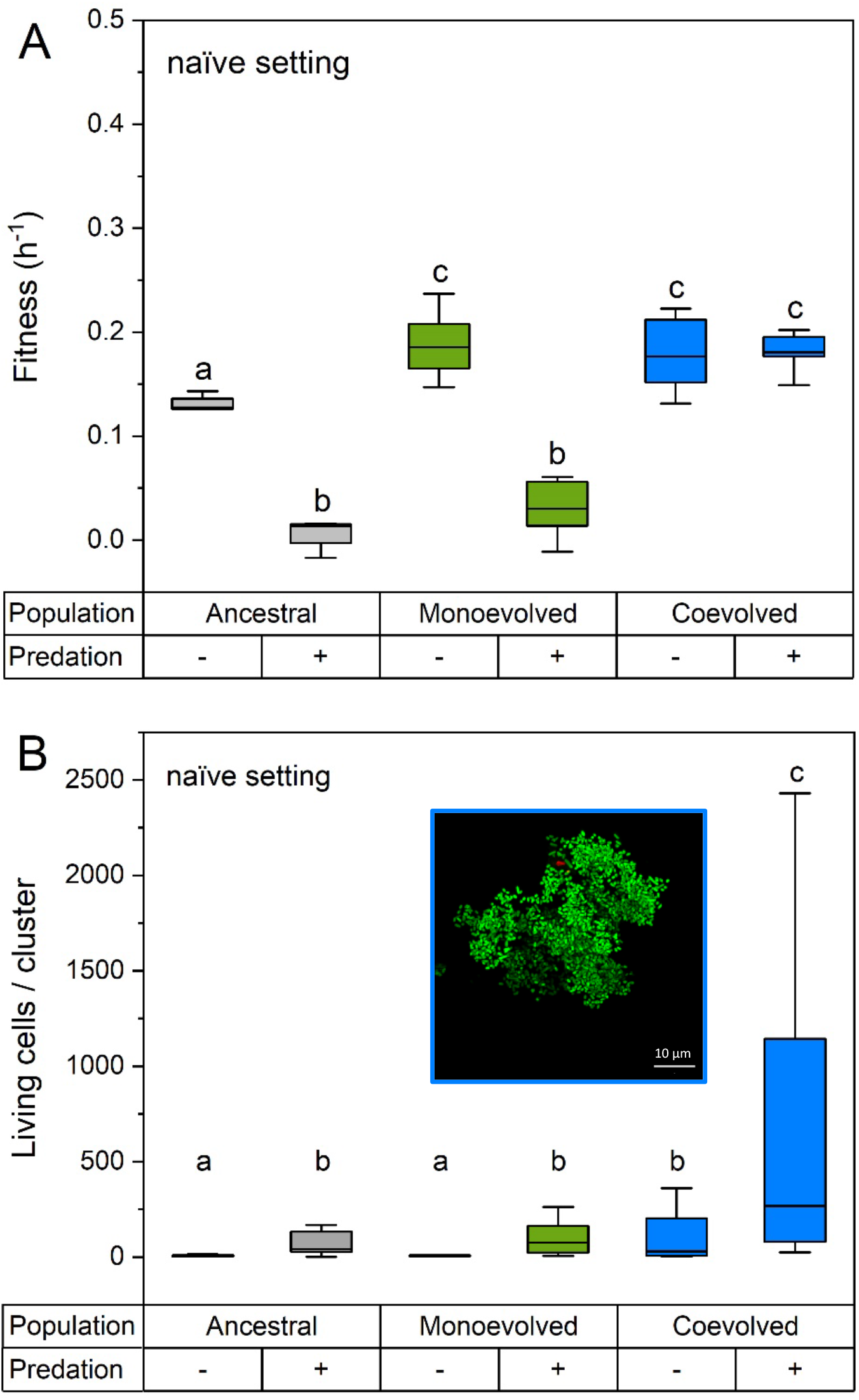
Under naïve conditions, predator-exposed offspring of coevolved populations shows high fitness and adaptive cluster formation. **(A)** Fitness and **(B)** cluster sizes of ancestral, monoevolved and coevolved cultures of *E. coli* in the absence (−) or presence (+) of the predator *T. thermophila* under naïve conditions (Figure 1B). Fitness is expressed as the Malthusian parameter for a growth period of 24 hours and cluster size as the number of living cells per cluster after growing for 24 hours. Different letters indicate significant differences in fitness (A: ANOVA followed by Tamhane’s T2 posthoc test of ln-transformed, absolute data: P < 0.001, n = 8) and cluster size (B: ANOVA followed by Fisher’s LSD posthoc test of ln-transformed data: P < 0.001, n = 12). Insert in (B) shows a representative cluster of the coevolved offspring in the presence of predation stained with a bacterial viability kit: green cells are alive (SYTO 9); red cells are dead (propidium iodide).

However, which ecological mechanism led to the observed increased protection of coevolved populations from *Tetrahymena* predation? Here we hypothesized that predation has induced the formation of multicellular aggregates, whose size exceeds the maximum capacity of *Tetrahymena* to ingest prey particles, thereby protecting cell clusters from predation. To verify this hypothesis, the same experimental populations that have previously been used to analyse the fitness consequences of predation (Figure 2A), were subjected to a microscopic analysis in order to quantify their propensity to form multicellular clusters. The results of this analysis confirmed indeed that in all three experimental groups (i.e. ancestral, monoevolved, and coevolved), predation resulted in the formation of significantly larger cellular clusters than were observed in the no-predator controls (ANOVA followed by Fisher’s LSD posthoc test of ln-transformed data: P_ancestral_ = 0.003, n = 12; P_monoevolved_ = 0.003, n = 12; P_coevolved_ = 0.001, n = 12, Figure 2B). Notably, the cluster sizes that the offspring of predator-free, coevolved populations formed was not significantly different from the ones the ancestral and monoevolved populations formed in the presence of predation (ANOVA followed by Fisher’s LSD posthoc test of ln-transformed data: both P > 0.3, n = 12, Figure 2B). Strikingly, the cluster size of coevolved offspring increased to a maximum of 2,430 living cells per single cluster in the presence of predation, thereby significantly exceeding the size of clusters observed in any of the other cultures (ANOVA followed by Fisher’s LSD posthoc test of ln-transformed data: all P < 0.012, n = 12, Figure 2B). Taken together, these results demonstrate that predation induces the adaptive formation of multicellular clusters and that this response is most pronounced in the offspring of coevolved populations.

### Predation increases fitness via the formation of multicellular clusters

Given that in the naïve setting predation decreased cellular fitness in monoevolved populations, but not in those that coevolved together with the predator, we were curious to find out how a temporally extended predation pressure would affect fitness and cluster formation in monoevolved versus coevolved populations. When cultures were grown in the presence of predators for three days (i.e. experienced setting), their fitness increased significantly over that of no-predator controls (ANOVA followed by Tamhane’s T2 posthoc test of square root-transformed data: P_monoevolved_ < 0.001, n = 8; P_coevolved_ = 0.001, n = 8, Figure 3A). This was not only true for the offspring of coevolved populations, but also for the monoevolved offspring, which had never experienced predation prior to these three days. These observations point to a secondary mechanism that is induced after three days and which positively effects the fitness of the respective cultures despite predation.

**Figure 3.**
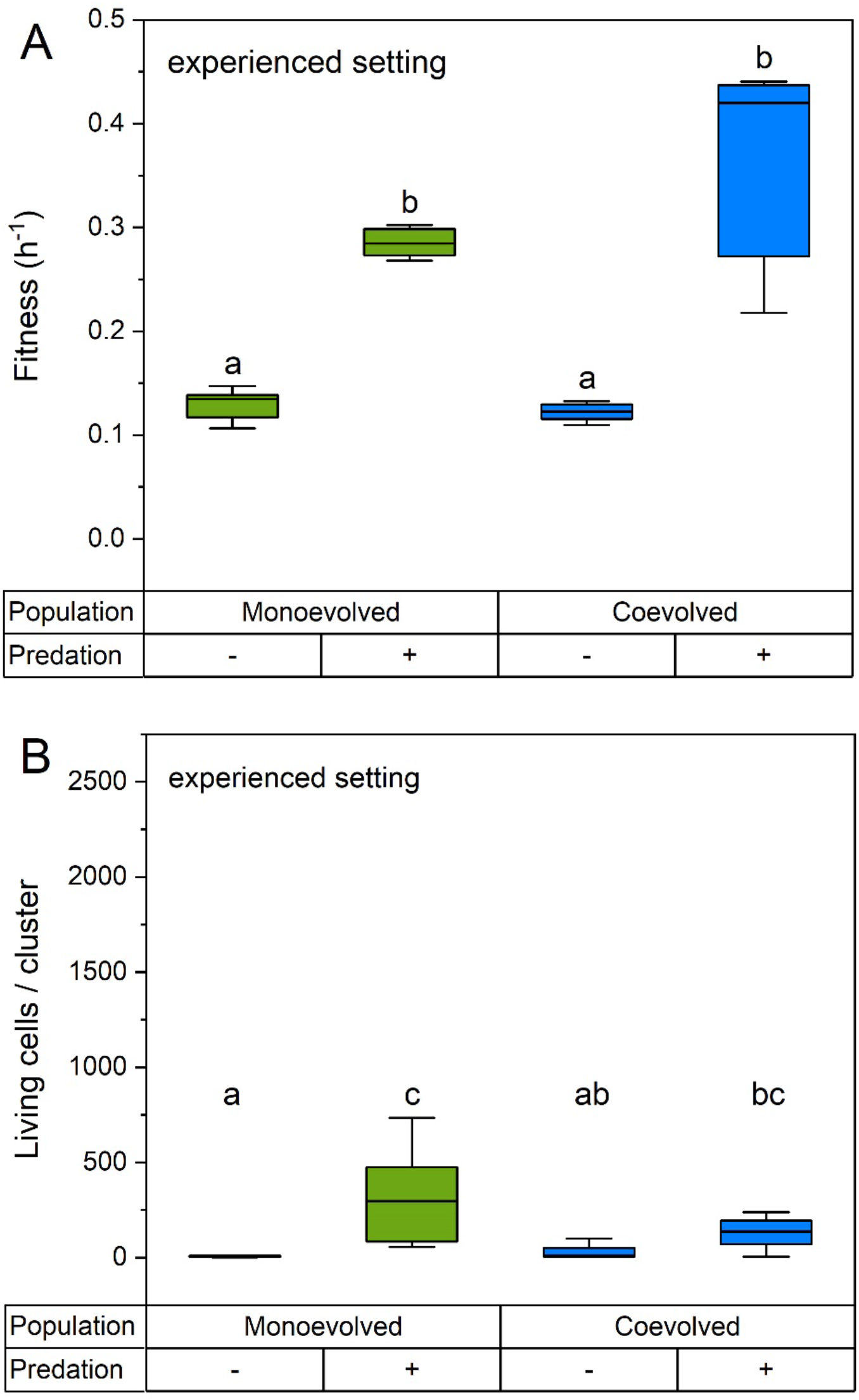
Under experienced conditions, predator-exposed offspring of mono- and coevolved populations shows an increased fitness despite small clusters. **(A)** Fitness and **(B)** cluster sizes of mono- and coevolved *E. coli* cultures in the absence (−) or presence (+) of the predator *T. thermophila* after three days of experiencing the respective condition (Figure 1C). Fitness is expressed as the Malthusian parameter for a growth period of 24 hours and cluster size as the number of living cells per cluster after growing for 24 hours. Different letters indicate significant differences in fitness (A: ANOVA followed by Tamhane’s T2 posthoc test of square root-transformed data: P < 0.001, n = 8) and cluster size of experienced populations (B: Kruskal-Wallis followed by Dunn’s multiple comparison test: P < 0.001, n = 12).

While clusters formed by populations of monoevolved cells were significantly larger in the presence of predation as compared to the no-predator control, the respective difference was less pronounced in offspring of coevolved cultures (Kruskal-Wallis followed by Dunn’s multiple comparison test: P_monoevolved_ < 0.001, n = 12; P_coevolved_ = 0.113, n = 12, Figure 3B). Nevertheless, in the face of predation, the size of clusters formed by monoevolved and coevolved populations did not differ significantly from each other (Kruskal-Wallis followed by Dunn’s multiple comparison test: P = 1.0, n = 12, Figure 3B). Together, these results support the idea of a secondary yet unknown effect playing a role that is independent of pure cluster size.

## DISCUSSION

Preparing your offspring for future challenges has significant consequences for the evolutionary fate of the entire population as it not only affects the strength and direction of natural selection, but also immediately enhances the offspring’s fitness. Thus, this mechanism appears to be particularly important in the context of predator-prey interactions. So far, however, evidence for transgenerational priming has been limited to eukaryotic organisms. Here we tested the hypothesis that predation primes bacterial populations such that their future generations can respond with a more effective induction of their defence mechanisms.

Our results show that in contrast to ancestral and monoevolved populations, populations that evolved in the presence of predators, became more resistant to predation (Figure 2A). This effect could be attributed to the formation of multicellular clusters (Figure 2B). The fact that our experiments were performed with the offspring of the respective cultures that was revived after freezing, demonstrates a transgenerational effect positively affected the fitness of coevolved populations. Thus, our data highlight the capability of bacteria to store and maintain information about ancestral stressful conditions, indicating bacterial memory across generations. Bacterial memory within individual cells has recently been demonstrated by Yang *et al*. (2020), who showed that blue light induces membrane-potential-based memory in single *Bacillus subtilis* cells residing within a biofilm. Moreover, *Pseudomonas aeruginosa* was reported to possess within- and between-generational memory as cells, which were previously exposed to a surface as well as the offspring of these cells, showed a stronger attachment to these surfaces later on (Lee *et al.*, 2018). Memory of acquired information together with the ability to use it in the future is key to the process of learning (Kawecki, 2010). More precisely, in learning, the phenotype is considered to depend not only on the genotype and the current environment, but also on the memory of past events. If the learned behaviour translates into improved fitness, learning becomes adaptive. An elegant example in this context is the grasshopper *Schistocerca americana* that learned to associate high food quality with certain cues like colour and flavour, thus experiencing higher growth rates than individuals prevented from employing associative learning (Dukas & Bernay, 2000). Interestingly, our data seem to be in line with the concept of adaptive learning as (i) the observed clustering phenotype not only depended on the current presence of predators, but also on the memory of past predation, and (ii) clustering enhanced the fitness of the respective populations. However, a learned response is generally regarded to develop within the lifetime of an individual and is usually based on sensory feedback (Kawecki, 2010). Whether the given framework should be extended to adaptive transgenerational learning in bacteria and which mechanisms might be involved to store and transmit information across generations, needs to be addressed in the future.

Besides adaptive learning, our cluster data meet the key criteria of priming. Comparing the cluster sizes of all three predator-exposed groups (Figure 2B), the offspring of coevolved populations showed the largest clusters. This indicates that offspring of populations that evolved in the presence of predators responded stronger to a current predation pressure than the offspring of ancestral and monoevolved cultures. This finding clearly points towards transgenerational priming and can explain how coevolved offspring manages to retain high fitness levels despite a prevailing predation pressure (Figure 2A). The observation that predator-free offspring of coevolved populations produced clusters that were comparable in size to the ones formed by predator-exposed offspring of ancestral and monoevolved cultures suggests transgenerational plasticity. However, clusters of coevolved offspring did not result in additional fitness advantages in the absence of predators. Moreover, ancestral and monoevolved cultures did not seem to benefit from the formation of predation-induced clusters (Figure 2A). Two factors are conceivable that could explain this pattern. First, the respective clusters might simply be too small to prevent them from being eaten by predators. Second, not only the cluster size and its defensive impact are crucial for the fitness of a given culture, but additionally a secondary cluster-related effect might play a role. If both, evolutionary history and time to interact, affect the manifestation of such an effect, this could explain the higher fitness of coevolved offspring as compared to the one of ancestral and monoevolved cultures. A potential explanation for this could be the phenomenon of division of labour. Indeed, division of labour has been demonstrated to readily evolve among cells of *Saccharomyces cerevisiae* that have been experimentally selected to form multicellular clusters (Ratcliff *et al.*, 2012). Interestingly, in our experiments, we identified two different morphotypes that mainly occurred in the coevolved offspring when exposed to predation. Unfortunately, these morphotypes turned out to be only transiently detectable and could therefore not be isolated and conserved to perform additional experiments.

However, the division of labour hypothesis bears the potential to explain observations in the experienced setting (Figure 3). After three days of continuous exposure to predation, the fitness of mono- and coevolved offspring significantly exceeded levels of the corresponding controls, while the respective clusters were not particularly large. In fact, the size of predation-induced clusters seemed to resemble those of the ancestral and monoevolved cultures from the naïve setting (Figure 2A). Since similar cluster sizes appeared to be correlated with differing fitness outcomes in the two different experimental settings, these comparisons support the idea of a cluster-associated division of labour, which, after it has developed, can positively affect the fitness of populations. Concerning the cluster size of coevolved offspring, one might ask why the transgenerational priming effect seen in the naïve setting cannot be observed in the experienced setting. The most plausible answer is that under continuous predation pressure, clusters evolve towards an optimal size that prevents them from being eaten and simultaneously allows for an efficient division of labour. Intriguingly, only three days of continuous predation seem to be sufficient to reach this optimum, as the monoevolved offspring, which has never been exposed to predators prior to the experiment, was characterized by an equally high fitness as the coevolved offspring (Figure 3A).

Taken together, our results demonstrate that bacteria under predation pressure are capable of transgenerational priming and that this phenomenon is more pronounced in the naïve setting than under the rather continuous conditions of the experienced setting. The latter observation is in line with the basic concepts of phenotypic plasticity and priming as previously described for eukaryotic organisms. In this context, phenotypic plasticity and inducibility are considered to be stress responses on demand (Karban & Baldwin, 1997; Schaller, 2008) and are thus more important in fluctuating environments than under constantly stressful conditions. Primability has been shown to be an elegant means to bridge the time delay until an effective induced defence is mounted (Hilker *et al.*, 2016; Martinez-Medina *et al.*, 2016). In this way, defence priming can help to fine-tune defence levels in the face of predation pressure that fluctuates in magnitude and frequency over time. However, our study highlights a significant difference compared to most previously reported cases of defence priming in eukaryotes: in bacteria such as *E. coli,* defence priming appears to operate across generations. Thus, it seems to represent a population-level rather than a pure individual-level response, which is reasonable considering the short generation time of bacteria due to which environmental fluctuations are more likely to affect multiple generations. Moreover, the fact that bacteria prepare their offspring for future challenges adds another level to our newly emerging perception of bacteria: while evidence is mounting that bacteria do not function as separated, autonomous units, but rather exist and operate within collectives (Pande & Kost, 2017), our study expands this view towards bacteria actively shaping their evolutionary fate.

## ACKNOWLEDGEMENTS

We thank Dr. Shraddha Shitut for the *Tetrahymena* drawings and the entire Kostlab (present and past) for useful discussions. This work was financially supported the University of Osnabrück (SK, LKM, and CK) and the international research training school *EvoCell* (LKM and CK).

